# Resource availability and disturbance shape maximum tree height across the Amazon

**DOI:** 10.1101/2020.05.15.097683

**Authors:** Eric Gorgens, Matheus Henrique Nunes, Tobias Jackson, David Coomes, Michael Keller, Cristiano Rodrigues Reis, Rubén Valbuena, Jacqueline Rosette, Danilo Roberti Alves de Almeida, Bruno Gimenez, Roberta Cantinho, Alline Zagnolli Motta, Mauro Assis, Francisca Rocha de Souza Pereira, Gustavo Spanner, Niro Higuchi, Jean Pierre Ometto

## Abstract

The factors shaping the distribution of giant tropical trees are poorly understood, despite its importance as a link between evolutionary biology and ecosystem biogeochemistry. The recent discovery of clusters of trees over 80 metres tall in the Guiana Shield region of the Amazon rainforest challenges the current understanding of the factors controlling the growth and survival of giant trees. The new discovery led us to revisit the question: what determines the distribution of the tallest trees of the Amazon?

Here, we used high-resolution airborne LiDAR (Light Detection and Ranging) surveys to measure canopy height across 282,750 ha of primary old-growth and secondary forests throughout the entire Brazilian Amazon to investigate the relationship between the occurrence of giant trees and the environmental factors that influence their growth and survival. Our results suggest that the factors controlling where trees grow extremely tall are distinct from those controlling their longevity. Trees grow taller in areas with high soil clay content (> 42%), lower radiation (< 130 clear days per year) and wind speeds, avoiding alluvial areas (elevations higher than 40 m a.s.l), and with an optimal precipitation range of 1,500 to 2,500 mm yr-1. We then used an envelope model to determine the environmental conditions that support the very tallest trees (i.e. over 70 m height). We found that, as opposed to the myriad of interacting factors that control the maximum height at a large scale, wind speed had by far the largest influence on the distribution of these sentinel trees, and explained 67% of the probability of finding trees over 70 m in the Brazilian Amazon forest.

The high-resolution pan-Amazon LiDAR data showed that environmental variables that drive growth in height are fundamentally different from environmental variables that support their survival. While precipitation and temperature seem to have lower importance for their survival than expected from previous studies, changes in wind and radiation regimes could reshape our forested biomes. This should be carefully considered by policy-makers when identifying important hotspots for the conservation of biodiversity in the Amazon.

## Introduction

The Amazon is the largest rain forest on Earth, covering 5.5 million square kilometres, and storing about 17% of all vegetation carbon. Ecologists have long taken an interest in comparing the structure and composition of rain forests across the tropics (Yang *et al*. 2016), and have reached a consensus that the Amazon supports shorter trees, and therefore stores a lower amount of carbon per hectare, than the forests of tropical Africa and Asia (Cao & Woodward 1998; Feldpausch *et al*. 2012). However, the recent discovery of giant trees - up to 88 m tall - in the Amazon basin (Gorgens *et al*. 2019) challenges this paradigm and poses new questions about the drivers causing the spatial distribution of tall trees in the Amazon.

Previous studies have debated the factors which govern Amazon tree growth and have particularly focused on productivity drivers related to the wet and dry seasons (Huete *et al*. 2006; Morton *et al*. 2014). This paper’s findings inform this important question but extend the investigation beyond these factors to include the influence of 18 climatic and other environmental conditions on achieving greatest tree height.

Tree height is fundamentally linked to growth, survival, and reproduction strategies, and is ultimately related to the ability to pre-empt light resources and disperse diaspores (Díaz *et al*. 2016). Xylem conduit diameter and total path length resistance to water flow increase with canopy height, making water transport to higher leaves more difficult (Koch *et al*. 2004; Givnish *et al*. 2014). To counteract this difficulty, taller species have higher xylem hydraulic conductivity but are more vulnerable to xylem embolism (Liu *et al*. 2019). Across species, higher wood density, and stomata closure in response to water deficit are often positively related to embolism resistance (Bennett *et al. 2015;* McDowell & Allen 2015; Greenwood *et al*. 2017). Height growth is partly governed by small-scale factors such as water availability, temperature, rooting depth, and soil type (Anderegg *et al*. 2016; McDowell & Allen 2015; Coomes *et al*. 2006; Niklas 2007), with precipitation and potential evapotranspiration consistently reported as key factors determining plant height across biomes (Moles *et al*. 2009; Larjavaara 2013; Rueda *et al*. 2016).

Forest giants are disproportionately vulnerable to disturbances and thus their conservation requires particular attention (Pennisi 2019; Yanoviak *et al*. 2019; Stovall *et al*. 2019; Enquist *et al*. 2020). To reach such immense sizes, trees must fulfill at least three conditions: they must (1) have an evolutionary design that is capable of transporting water to great heights and overcome highly negative water potentials to deliver that water toward tissues in the upper canopy; (2) inhabit an area with optimal environmental conditions (such as climate, soil properties, and water) that meet species-specific requirements (Simard *et al*. 2018; Scheffer *et al*. 2018) and (3) grow in regions with a low frequency of natural or anthropogenic disturbance events (Larjavaara 2013; Linden-mayer & Laurance 2016; Scheffer *et al*. 2018; Enquist *et al*. 2020). Resource availability (e.g. sunlight, nutrients, CO_2_, and water) controls a tree’s ability to produce biomass through photosynthesis. Natural disturbances (e.g. windthrow, drought, or lightning) and history of anthropogenic actions (e.g. selective logging, forest fragmentation) increase the likelihood of mortality and limit the time available to trees to grow taller (Bennett *et al*. 2015; Powers *et al*. 2020; Yanoviak *et al. 2019*; Almeida *et al*. 2019). Tall trees are likely to have developed strategies for surviving diseases and pathogens (van Gelder *et al*. 2006; Aleixo *et al*. 2019) as well as climatic fluctuations (Sakschewski *et al*. 2016) and resisting wind damage (Jagels *et al*. 2018). However, the question of how resource supply and disturbances interact to determine canopy height across the Amazon has not been fully explored.

The sheer size of the Amazon, its environmental heterogeneity and species diversity, pose challenges and practical difficulties to understand general ecological relationships and biogeographical patterns (Tuomisto *et al*. 2019). Forest plots provide many valuable insights to investigate the influences of the environment on tree height but they can only represent a minuscule fraction of the total forest area (Chave *et al*. 2020). Currently, a network of 5,351 forest inventory plots established across the Brazilian Amazon, of known and published sites recently compiled by (Tejada *et al*. 2019), represents only 0.0013% of the total forest area in this region. In addition, the plot distribution is spatially clustered in close proximity to major roads or large rivers (Stropp *et al*. 2020), implying a spatial distribution bias (Marvin *et al*. 2014) since about 42% of the total Brazilian Amazon lies over 50 km from the nearest forest inventory plots (Tejada *et al*. 2019). Remote sensing can remove sampling biases and uncertainty about ecological patterns (Schimel *et al*. 2015) and provides large datasets with which to uncover the environment controls of forest structure (Asner *et al*. 2010). In particular airborne LiDAR (Light Detection and Ranging) generates valuable high-resolution 3D information of forest canopy structure (Görgens *et al*. 2016; Coomes *et al*. 2017), and can be used as an intermediary to integrate field data with satellite sources (Asner 2009; Bae *et al*. 2019).

The “Improving Biomass Estimation Methods for the Amazon” (EBA) project performed high-resolution LiDAR flights over 3 years, totaling more than 800 transects across mature and secondary forests in the Brazilian Amazon (for more information about the EBA project see the Method section). The transects were randomly distributed considering both spatial location of the start point and flight direction, allowing us to conduct statistical design-based models of tree height since there is a growing consensus that ecosystem traits like tree height can be better measured with LiDAR than other methods (Valbuena *et al*. 2020). A total of 282,750 ha were covered, 0.183% of the Brazilian Amazon, which is 100 times the area of available permanent plot networks (Tejada *et al*. 2019). This unprecedented dataset has led to remarkable discoveries in Amazon (Gorgens *et al. 2019*; Pereira *et al*. 2019; Santos *et al*. 2019; Almeida *et al*. 2019). In this study, we employed it to contribute to our understanding of how resources and disturbances shape the maximum height distribution across the Brazilian Amazon. We conducted an extensive analysis relating remotely sensed environmental variables to the maximum height recorded in the transects. We concluded that drivers of height development are fundamentally different from those influencing the survival of tree giants. Thus changes in wind and light availability shape their distribution as much as precipitation and temperature, altogether shaping the demographics and composition of forested biomes.

## Material and methods

Between 2016 and 2018, an airborne mission (held by National Institute for Space Research - INPE and funded by Amazon Fund) collected airborne LiDAR data from 906 transects of 375 ha (12.5 x 0.3 km) each, randomly spread across primary and secondary forests defined by the PRODES database - layer mask of primary old-growth forests (**PRODES, INPE**, 2016) and by the TerraClass database - a layer mask of secondary forest (**TerraClass, INPE**, 2014).

The LiDAR sensor was the Trimble Harrier 68i (Trimble, California, USA) aboard a Cessna 206 aircraft. The average pulse density was set at 4 pulses m^-2^, the field of view equal to 30°, and flying altitude of 600 m. The Global Navigation Satellite System (GNSS) collected data on a dual-frequency receiver (L1/L2). The pulse footprint was set to be below 30 cm, based on a divergence angle between 0.1 and 0.3 mrad. Horizontal accuracy was controlled to be under 1 m, and the vertical accuracy to be under 0.5 m.

Details about LiDAR parameterization, processing, and the EBA project characteristics can be consulted in (Gorgens *et al*. 2019). Briefly, each transect was processed by identifying the returns backscattered from the ground and interpolating a 1m spatial resolution digital terrain model (DTM) from them. Then, the DTM was employed to calculate the heights above ground from the returns backscattered from the vegetation (Görgens *et al*. 2016). The uppermost vegetation heights were then employed to compute a canopy height model CHM at the same spatial resolution as the DTM. The height of the tallest tree per transect was identified from the CHM using a local maximum moving window algorithm (Dalponte & Coomes 2016). All transects were finally manually inspected to exclude non-trees maximum derived from artifacts, ensuring that all the largest heights indeed depicted a tall tree.

### Environmental variables

To investigate drivers influencing the spatial distribution of giant trees, we initially considered a total of 18 environmental variables: (1) fraction of absorbed photosynthetically active radiation (FAPAR; in %); (2) elevation above sea level (Elevation; in m); (3) the component of the horizontal wind towards east, i.e. zonal velocity (u-speed; in m s^-1^); (4) the component of the horizontal wind towards north, i.e. meridional velocity (v-speed; in m s^-1^); (5) the number of days not affected by cloud cover (clear days; in days yr^-1^); (6) the number of days with precipitation above 20 mm (days > 20mm; in days yr^-1^); (7) the number of months with precipitation below 100 mm (months < 100mm; in months yr^-1^); (8) lightning frequency (flashes rate); (9) annual precipitation (in mm); (10) potential evapotranspiration (in mm); (11) coefficient of variation of precipitation (precipitation seasonality; in %); (12) amount of precipitation on the wettest month (precip. wettest; in mm); (13) amount of precipitation on the driest month (precip. driest; in mm); (14) mean annual temperature (in °C); (15) standard deviation of temperature (temp. seasonality; in °C); (16) annual maximum temperature (in °C); (17) soil clay content (in %); and (18) soil water content (in %).

The FAPAR was derived from land surface reflectance product calibrated and corrected from the National Oceanic and Atmospheric Administration’s (NOAA) Advanced Very High-Resolution Radiometer (AVHRR), which is a consistent time-series dataset spanning from the mid-1980s to present and suitable for climate studies (Tao *et al*. 2016). FAPAR is a primary vegetation variable controlling the photosynthetic activity of plants and is considered an essential climate variable (Mason *et al*. 2010). The algorithm to create this layer relies on artificial neural networks calibrated with the MODIS FAPAR dataset and validated using a set of globally-distributed sites. The inputs to generate the FAPAR were 1) the surface reflectance from NOAA-AVHRR which is measured in two wavelengths (red, 580–680 nm, and near-infrared, 725–1000 nm); 2) a reference dataset from MODIS FAPAR to calibrate the NOAA-AVHRR FAPAR; and 3) a land cover map classification, used to stratify the outputs. Five land cover classes were included: evergreen broadleaf forest, deciduous broadleaf forest, needle leaf forest, shrubland, croplands and grasslands, and non-vegetated (Claverie *et al*. 2016).

The elevation was computed based on the third version of the Shuttle Radar Topography Mission (SRTM) provided by National Aeronautics and Space Administration Jet Propulsion Lab (NASA JPL) (Farr *et al*. 2007; Liu *et al*. 2014). The SRTM mission was launched on Space Shuttle Endeavor on 11th February 2000 and collected data during ten days of operations, using two synthetic aperture radars: NASA’s C band system (5.6 cm wavelength) and an X band system by DLR (3.1 cm). The C-band digital elevation model (DEM) used in this study is now available at 30-m spatial resolution from 60° north latitude and 56° south latitude, covering 80% of Earth’s land surface. C-band partially penetrates the vegetation canopy, with depth varying with vegetation structure. Since Amazonian vegetation is dense throughout, for the purposes of this study the C-band DEM is assumed to vary consistently with topography across the region.

We used the maximum daily mean wind speeds over the last 5 years from the fifth major global reanalysis (ERA5) produced by the European Centre for Medium-Range Weather Forecasts (ECMWF). The reanalysis combined model data with observations from across the world into a globally complete and consistent dataset (Olauson 2018). The products from a reanalysis include many variables such as wind speeds, temperature, and atmospheric pressure. They were produced on reduced Gaussian grids, by using a different number of grid points along different latitudes and thus keeping the grid point separation in metres approximately constant. ERA5 has an hourly resolution and spans from 1950 to near real-time. Two wind velocities were considered: u-speed which is the zonal velocity (i.e. the component of the horizontal wind towards east), and v-speed which is the meridional velocity (i.e. the component of the horizontal wind towards north). These products are used extensively for modelling wind power both in academia and industry (Olauson 2018; Albergel *et al*. 2019; Ramon *et al*. 2019).

The number of clear days was computed based on Moderate Resolution Imaging Spectroradiometer (MODIS) surface reflectance products. MODIS products provide an estimate of the surface spectral reflectance as it would be measured at ground level in the absence of atmospheric scattering or absorption (Kang *et al*. 2005; Bisht & Bras 2010). MODIS operates onboard Terra and Aqua satellites. Terra satellite has a 10:30 am equator over-passing time, and the ±55° scanning pattern at 705 km altitude achieves a 2,330 km swath that provides global coverage every one to two days (Ruhoff *et al*. 2012). We used the Terra MOD09GA Version 6 product, which provides an estimate of the surface spectral reflectance of MODIS, corrected for atmospheric conditions such as gases, aerosols, and Rayleigh scattering.

Temperature and precipitation were obtained from the WorldClim database of bioclimatic variables, which are derived from weather station data compiled for the 1950-2000 period (Hijmans *et al*. 2005; Fick & Hijmans 2017). The main source of data was the Global Historical Climatology Network (GHCN), complemented with other global, national, regional, and local data sources, which were added if they were further than 5 km away from stations already included in the GHCN. After removing stations with errors, the final database consisted of precipitation records from 47,554 locations, mean temperature from 24,542 locations, and minimum and maximum temperatures from 14,835 locations. To interpolate the weather station data, latitude, longitude, and elevation were used as independent variables.

The lightning frequency was provided by Lightning Imaging Sensor (LIS) instrument onboard the Tropical Rainfall Measuring Mission provided by NASA Earth Observing System Data and Information System (EOSDIS) Global Hydrology Resource Center. The LIS was launched in November 1997 into a precessing orbit inclination of 35° at an altitude of 350 km and was powered off in April 2015. The LIS datasets were collected during 16 years (1998–2013) and they are available at 0.1° spatial resolution (approx. 11km in the equator). The LIS provided the basis for the development of a comprehensive global thunderstorm and lightning climatology to detect the distribution and variability of total lightning occurring in the Earth. This information is used for severe storm detection and analysis, and also for lightning-atmosphere interaction studies (Albrecht *et al*. 2016).

The potential evapotranspiration was provided by the TerraClimate dataset, a global monthly climate and water balance for terrestrial surfaces spanning 1958–2015. The layer used climatically aided interpolation (bilinear interpolation of temporal anomalies), combining high-spatial-resolution climatological normals from WorldClim with Climate Research Unit (CRU) Ts4.0 and the Japanese 55-year Reanalysis (JRA-55) data. The CRU Ts4.0 provides monthly average maximum and minimum temperature, vapor pressure, and cumulative precipitation from 1901–2015. The JRA-55 is the longest-running (1958-present) observing-system and provides spatially and temporally complete data for mean temperature, vapor pressure, wind speed, downward shortwave flux at the surface, and accumulated monthly precipitation. The Reference Evapotranspiration was calculated using the Penman-Monteith approach (Abatzoglou *et al*. 2018).

The number of months per year with precipitation below 100 mm and the number of days per year with precipitation above 20 mm was computed based on the Climate Hazards Group InfraRed Precipitation with Station data (CHIRPS) dataset. CHIRPS incorporated 0.05° resolution satellite imagery with in-situ station data to create gridded rainfall time series for trend analysis and seasonal drought monitoring (Funk *et al*. 2015). The CHIRPS process involves three main components: 1) the Climate Hazards group Precipitation climatology (CHPclim), 2) the satellite-only Climate Hazards group Infrared Precipitation (CHIRP), and 3) the station blending procedure that produces the CHIRPS. Two sets of monthly historical long-term means were used to create the CHPclim. The first set was a collection of 27,453 monthly stations obtained from the Agromet Group of the Food and Agriculture Organization of the United Nations (FAO). The second set of 20,591 stations was taken from version two of the Global Historical Climate Network (GHCN). The CHIRP relies on two global thermal infrared archives that are: the 1981–2008 Globally Gridded Center Satellite (GriSat) produced by NOAA’s National Climate Data and the 2000-present dataset NOAA Climate Prediction Center. The CHIRPS station datasets were obtained from the GHCN monthly, GHCN daily, Global Summary of the Day, Global Telecommunication System and Southern African Science Service Centre for Climate Change and Adaptive Land Management. The station blending procedure that produces CHIRPS is a modified inverse distance weighting algorithm. Daily CHIRPS are then produced for the globe by using daily Cold Cloud Duration (CCD) data to identify non-precipitating days. Whenever the daily CCD is zero, precipitation is assumed to be zero.

Edaphic variables were obtained from The OpenLandMap produced by the OpenGeoHub Foundation and contributing organizations. Soil texture is connected with soil granulometry or the composition of the particle sizes (clay, silt, and sand), typically measured as volume percentages. The clay content (fine particles < 2 μm) and water content layers, both with a spatial resolution of 250 m, were created based on machine learning predictions from a global compilation of soil profiles and samples (Arsanjani *et al*. 2014).

To help visualization of the regional-level, we divided the Brazilian Amazon into eight regions, according to the classification of (Morrone 2014): I - Para; II - Xingu-Tapajos; III - Roraima; IV - Guianan Lowlands; V - Madeira; VI - Yungas; VII - Pantepui; VIII - Imeri. This regionalization is based on biogeographic analyses of terrestrial plant and animal taxa of the Neotropical region and seeks to provide universality, objectivity, and stability, such that it can be applied when describing distributional areas of particular taxa or comparing different biogeographic analyses.

### Random Forest and Maximum Entropy

To better understand the environmental requirements for development in tree height, we employed Random Forest modelling and marginal plots to observe the relative variable importance. Among the initial 18 environmental variables, two of them (precipitation on driest month and months < 100mm) were excluded due to high correlation (> 0.80) to other independent variables (Table 1). Using the coordinates of the tallest tree within each lidar transect, we extracted the values from the variable layers. Tree height was then modeled against the factors using a random forest algorithm, which recursively computes classification and regression trees (CART) from random subsets, a k-fold (k = 15) cross-validation method, and 500 as the number of CART. The number of variables randomly sampled as candidates at each split was set to 10. The adjusted model was evaluated considering the mean absolute error (MAE), root mean squared error (RMSE), and coefficient of determination (R^2^) of cross-validated predicted versus observed values. To assess the overall relative variable importance we used the mean decrease in Gini importance, which evaluates at each split in each tree, how much each variable contributes to decreasing the weighted impurity (i.e., variance in the case of regression trees). The resulting Random Forest model was finally implemented using the environmental variables to deliver a map of estimated heights of tallest trees across the Amazon. Then we focused on the tallest trees only - those over 70 m in height - to determine the conditions which allow them to survive. We employed a maximum entropy envelope approach (MaxEnt) commonly applied to modelling species geographic distributions with presence-only data and indicate better discrimination of suitable versus unsuitable areas for the species (Phillips *et al*. 2006). The variable importance of the MaxEnt model was used to indicate the most relevant characteristics sustaining extreme height individuals and the potential location for new occurrence. The observations higher than 75 m were filtered out and used to adjust an envelope model based on maximum entropy. In its optimization routine, the algorithm tracked how much the model gain was improved when small changes were made to each coefficient value associated with a particular variable. Each variable was then ranked based on the proportion of all contributions. The resulting MaxEnt model was finally implemented using the environmental variables to deliver a map of probability of occurrence for trees taller than 70 m across the Amazon.

**Table 1:**
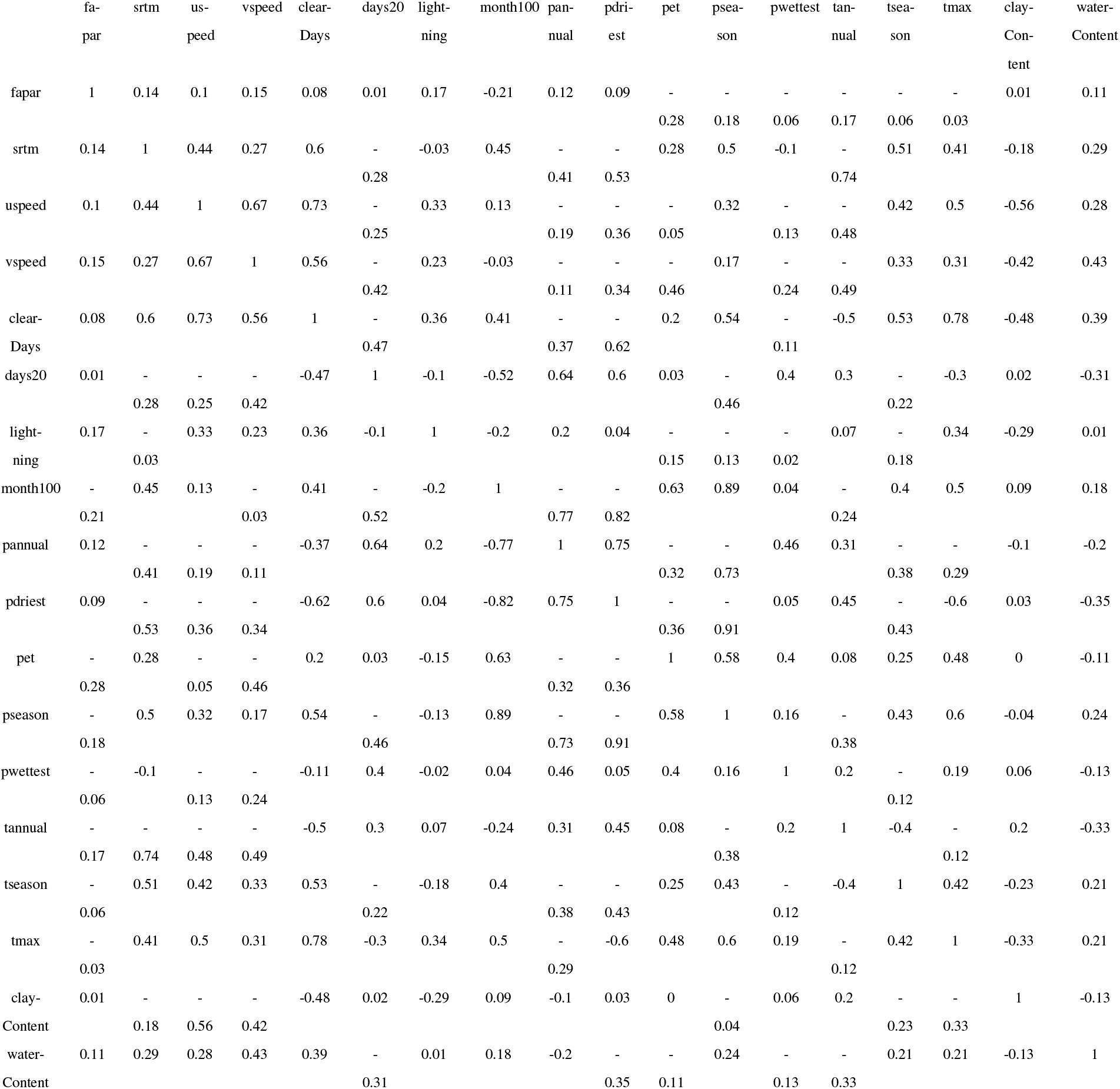
Correlation between the environmental variables.

## Results

Trees exceeded 50 m in height across many parts of the Brazilian Amazon, but trees over 80 m were only observed in the eastern Amazon (micro-region III, Roraima Province; Fig. 1). To examine why the trees grow taller in some regions and determine the environmental variables modulating height pattern in the Amazon, we predicted maximum tree height as a function of environmental variables using a Random Forest approach. The number of clear days, clay content in the soil, elevation and mean annual precipitation were found to be the strongest drivers of maximum tree height, while the average monthly temperature and soil water content were weak predictors (Table 2). The Random Forest model obtained MAE = 3.62 m, RMSE = 4.92 m, and R^2^ = 0.735. A resulting map of Random Forest model predicted maximum tree height shows that occurrence is highest in eastern Amazon (Fig. 2), with the tallest trees more specifically achieving greatest heights in the northeastern part of Roraima (III), in Pantepui (VII) and in the confluence of Madeira (V) and Xingu-Tapajos (II). Since low values of FAPAR are related to degraded forests and anthropogenic regions, we performed the same analysis after excluding areas with FAPAR values under 80%, which resulted in the elimination of 133 transects. Similar spatial distributions for maximum tree height persisted similarly after removing these potential anthropogenic effects (Fig. 3), demonstrating that the underlying patterns we report are naturally driven by the environmental factors.

**Figure 1.**
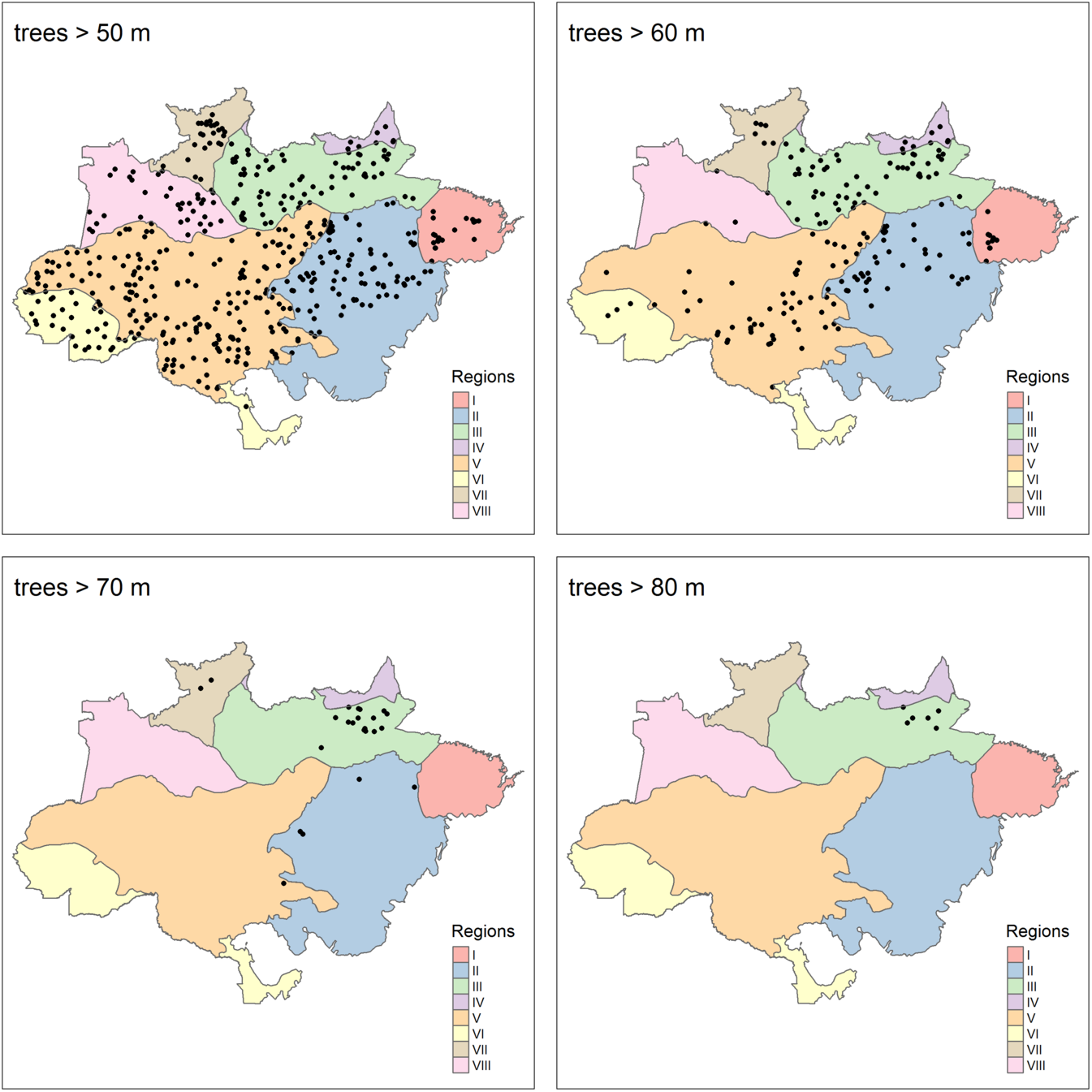
Maps of the Brazilian Amazon showing the location of trees > 50 m, > 60 m, > 70 m, and > 80 m in height. Black circles indicate the presence of a tree above the height thresholds. Background color indicates the biogeographical subdivisions proposed by (Morrone 2014): I - Para; II - Xingu-Tapajos; III - Roraima; IV - Guianan Lowlands; V - Madeira; VI - Yungas; VII - Pantepui; VIII - Imeri.

**Figure 2.**
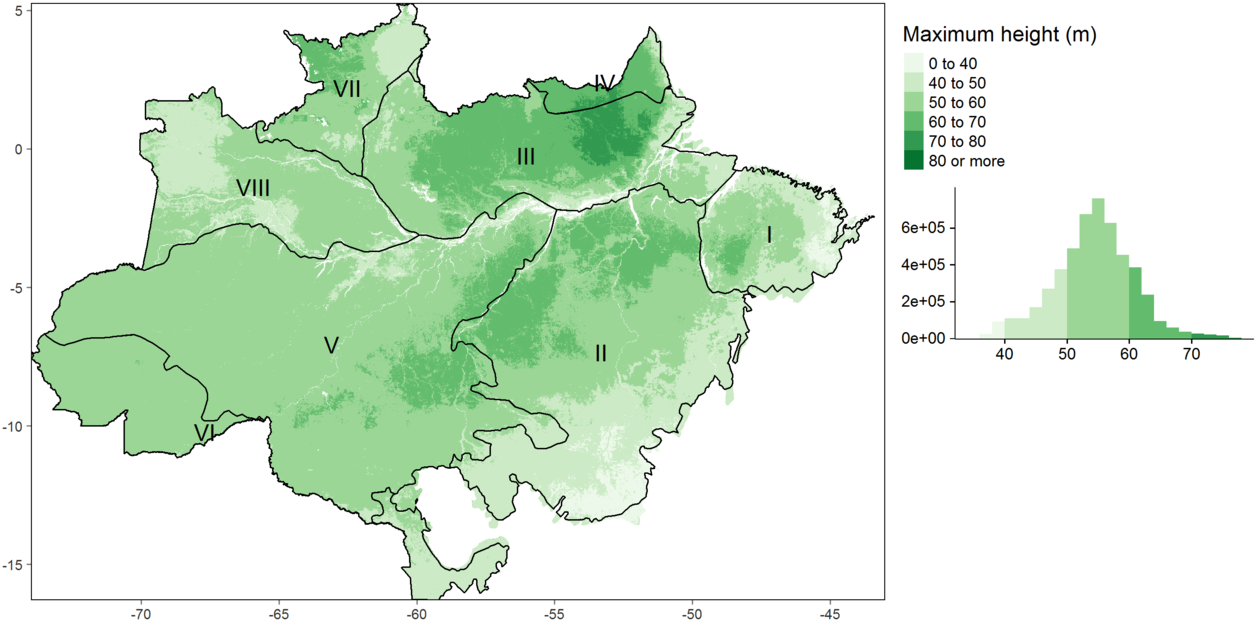
Maximum height estimation based on remote sensing variables estimated by Random Forest method. Black lines indicate the biogeographical subdivisions: I - Para; II - Xingu-Tapajos; III - Roraima; IV - Guianan Lowlands; V - Madeira; VI - Yungas; VII - Pantepui; VIII - Imeri.

**Figure 3.**
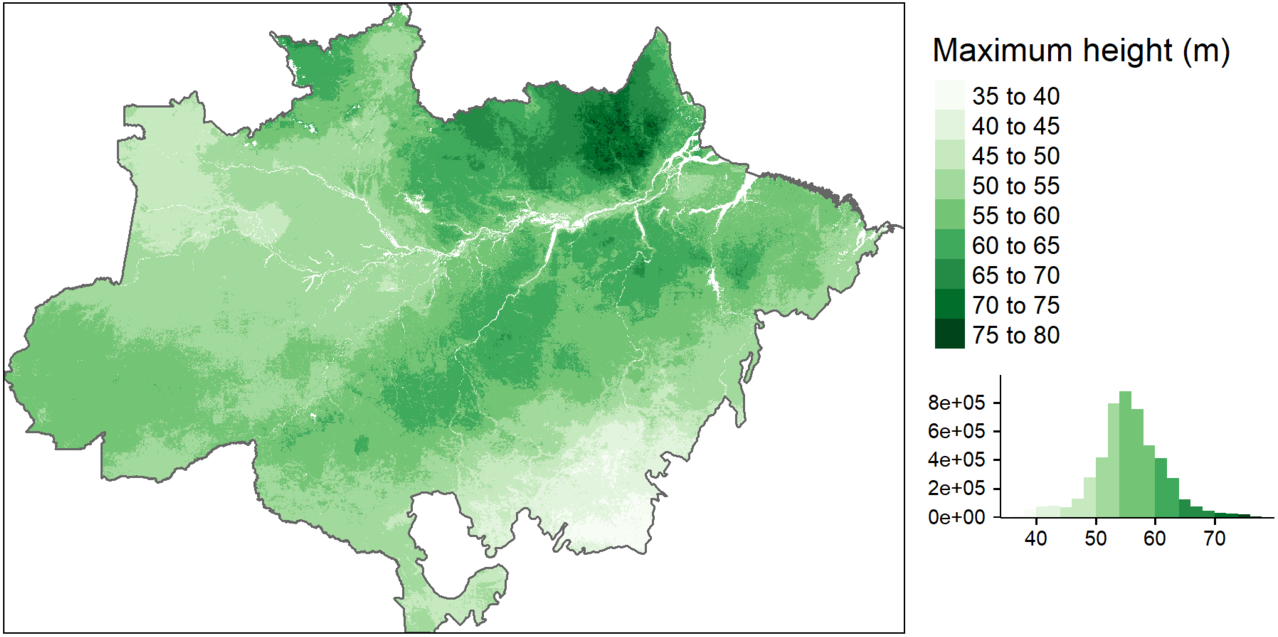
Maximum tree height distribution when FAPAR values under 80% were excluded from our analysis.

**Table 2:** Variables used to estimate maximum height distribution and evaluate its distribution, ranked by variable importance results in the Random Forest model

| Layer | Definition | Related to | Unit | Source | Importance |
| --- | --- | --- | --- | --- | --- |
| clearDays | number of clear days per year | energy balance - water balance - radiation | days | MODIS | 14.7 |
| clayContent | fraction of clay content | soil structure - physical properties - water availability | % | Open-LandMap | 13.8 |
| topography | elevation above sea level | distance to water - flooding zones - soil | m | SRTM | 11.2 |
| pannual | average annual precipitation | precipitation - precipitation intensity - precipitation distribution | mm | WorldClim | 8.9 |
| pseason | precipitation seasonality | precipitation - precipitation intensity - precipitation distribution | mm | WorldClim | 6.9 |
| fapar | fraction of absorbed photosynthetically active radiation | radiation - vegetation health - anthropic regions - soil exposure | % | NOAA AVHRR | 6.3 |
| pwettest | precipitation of the wettest month | precipitation - precipitation intensity - precipitation distribution | mm | WorldClim | 5.8 |
| uspeed | zonal speed (W-E) | storms - convective winds | m/s | ECM-RWF | 5.6 |
| days20 | days with precipitation higher then 20 mm | storms - convective winds | days | CHIRPS | 5.5 |
| pet | potential evapotranspiration | energy balance - water balance - radiation - vegetation health - anthropic regions - soil exposure | mm | TerraClimate | 5.2 |
| tseason | temperature seasonality | temperature - temperature distribution | C | WorldClim | 4.6 |
| tmax | maximum temperatura | storms - convective winds | C | WorldClim | 4.2 |
| vspeed | meridional speed (N-S) | storms - convective winds | m/s | ECM-RWF | 3.6 |
| lightning | lightning rate | storms - convective winds | flashes rate | LIS TRMM | 3.5 |
| tannual | daily average annual temperature | temperature - temperature distribution | C | WorldClim | 0.3 |
| water-Content | fraction of water content | soil structure - physical properties - water availability | % | Open-LandMap | 0 |
| month100 | month with precipitation below 100 mm | precipitation - precipitation intensity - precipitation distribution | months | CHIRPS | Removed by high correlation |
| pdriest | precipitation of the driest month | precipitation - precipitation intensity - precipitation distribution | mm | WorldClim | Removed by high correlation |

The marginal plot obtained for each environmental variable in the random forest model, allows us to interpret its influence on the height of tall trees directly in the units that correspond to each (Fig. 4). Lines close to horizontal indicate a given environmental factor having little effect on the height of tall trees. The number of clear days was the strongest predictor of maximum height (Table 2). The shape of this relationship resembles a step function (Fig. 4), in which regions with the number of clear days below 130 days per year support tall trees, above this level, we observe an abrupt decline in maximum height. Elevation was also a key predictor of tree height, with low-lying forests growing 7 m lower than trees in terrains above 40 m above sea level. An increase in soil clay content from 20% to 40% translated into a 7 m increase in maximum height. Our results also demonstrate mean annual precipitation as a key factor for trees to grow taller, with a tolerance curve peaking at around 2,300 mm yr^-1^ as optimal annual precipitation across the Brazilian Amazon. In comparison to these areas, we observe a 4 m decline in maximum tree height in regions with annual precipitation below 1,500 mm yr^-1^ or above 3,000 mm yr^-1^. From the intermediate importance variables, we highlight the zonal velocity (u-speed) and FPAR influencing height variation in ranges around 6 m.

**Figure 4.**
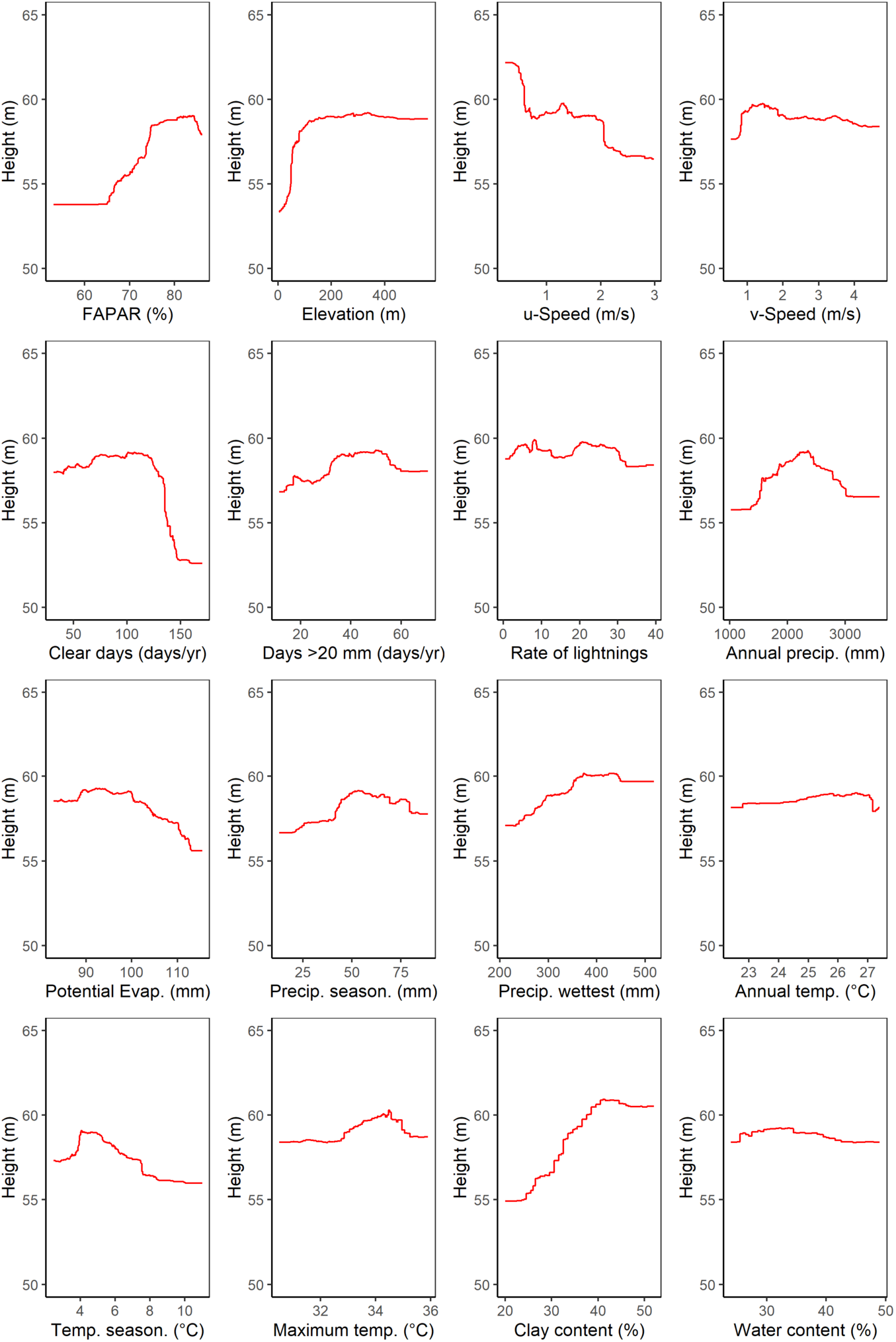
Marginal plot for each variable considering the Random Forest model for maximum height estimation

The results of the MaxEnt approach are focusing on the survival of trees taller than 70 m in height (Fig. 5). The extraordinarily tall trees had a unique niche, characterized by a much smaller set of environmental variables from those which drove the large-scale patterns of maximum height.

**Figure 5.**
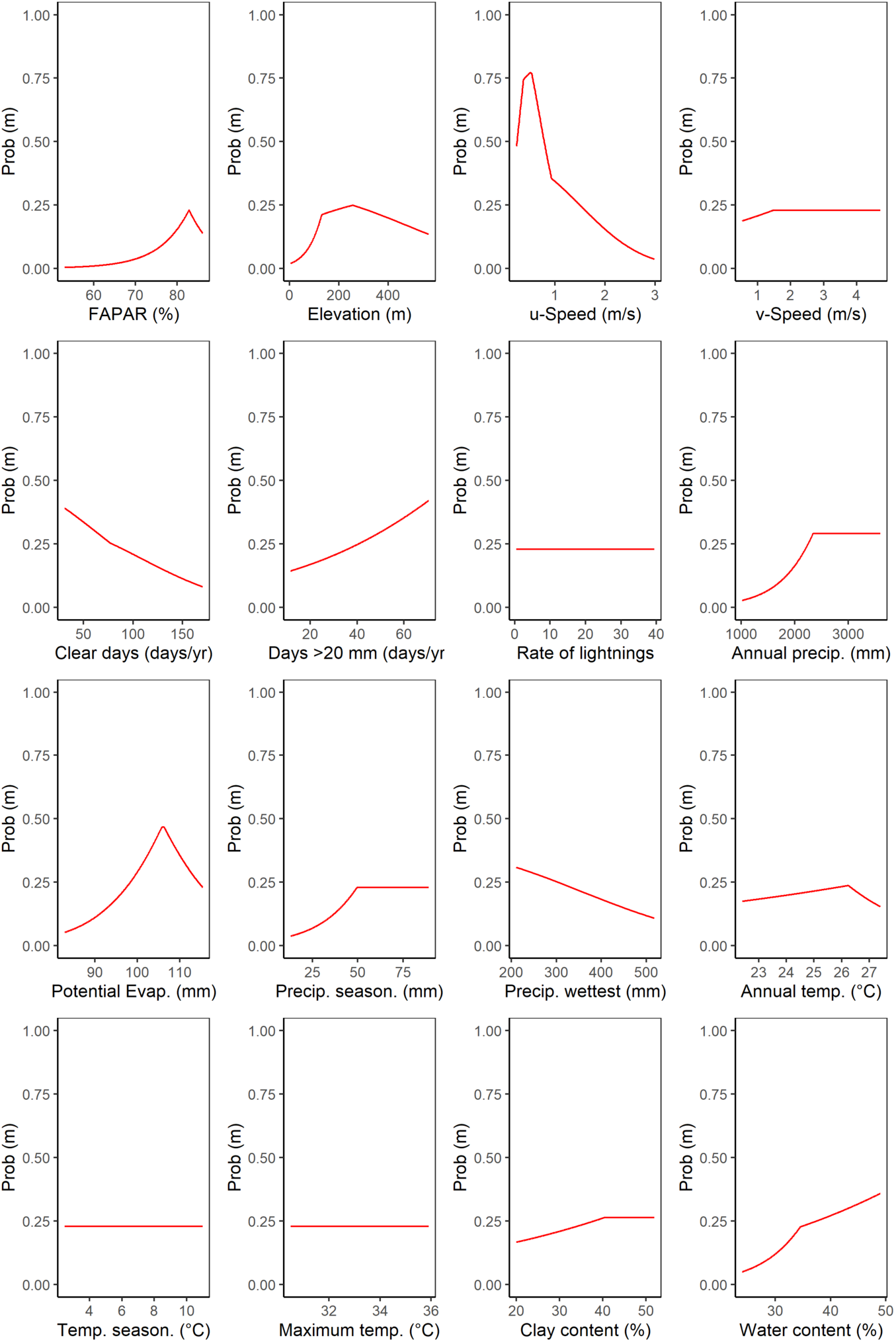
Marginal plot for each variable considering the Maximum Entropy model for niche determination

The maximum entropy model shows that the niche is dominated mostly by wind speed (relative importance of 67.7 %). The second most important driver of tall tree survival was the elevation above sea level (relative importance of 12.3 %). It is worthwhile noting that relative importance values reflect the proportion of all contributions to explain the presence of the tallest trees. The resulting map of predicted occurrence of the tallest trees in the Amazon from the MaxEnt model shows that the probability of maximum tree height occurrence is highest in northeastern Amazon (Fig. 6), more specifically in the Roraima (III) and Guianan Lowlands (IV).

**Figure 6.**
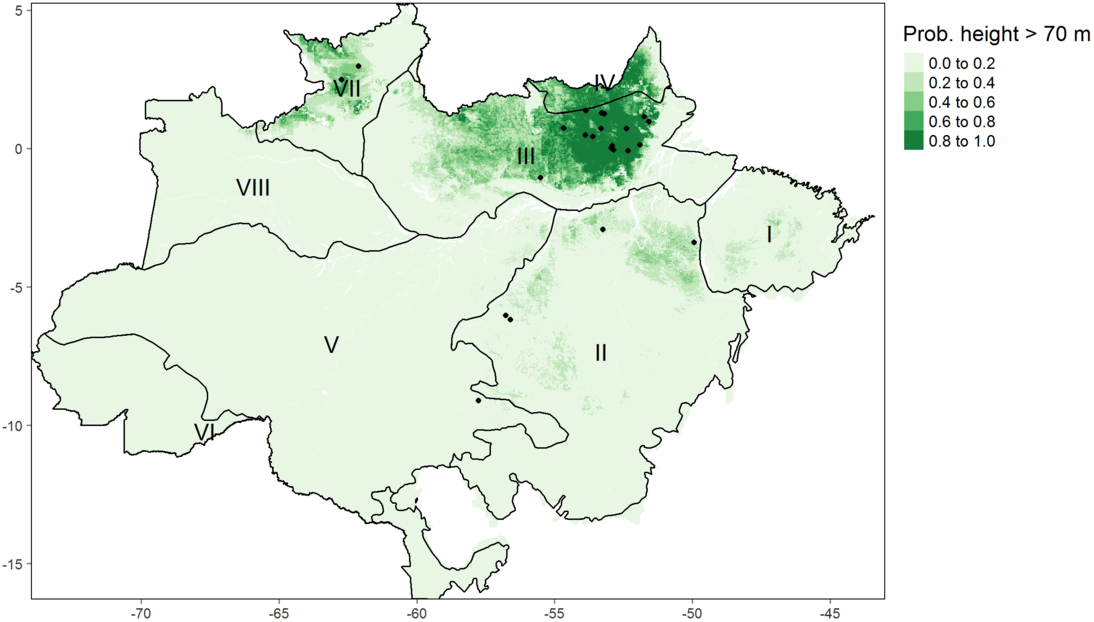
Ocurrence of giant trees (black dots) and niche capability to support the development of tall trees (probability of tall tree ocurrence). Black lines indicate the biogeographical subdivisions: I - Para; II - Xingu-Tapajos; III - Roraima; IV - Guianan Lowlands; V - Madeira; VI - Yungas; VII - Pantepui; VIII - Imeri.

## Discussion

The locations of the tall trees in the eastern and southern Amazon coincide with forests that have a high basal area predicted by statistical modelling of permanent plot data (Malhi *et al*. 2006). The basal area generally declines with increasing dry season length, for regions with dry seasons lasting four months or longer. Young soils nearer the Andes, as well as the sedimented and flooded lowlands, are richer in nutrients, thereby supporting fast-growing, low wood density species with high turnover rates and, as a result, the trees do not reach extremely large sizes (Marra *et al*. 2014; Quesada *et al*. 2011; Phillips *et al*. 2004). The species *Dinizia excelsa* (Ducke), for example, has been reported as the tallest trees in the Amazon reaching 88 m in height in the region of Roraima (Gorgens *et al*. 2019), and also has been reported as the highest average species-level wood density in the Amazon of 0.94 g cm^-3^ (Fauset *et al*. 2015) with a large contribution to the total forest biomass.

Many physiological and structural traits in the Amazon have strong phylogenetic associations with effects on tree growth and mortality (Baker *et al*. 2004; Fyllas *et al*. 2012; Patiño *et al*. 2012). Forests of the western Amazon are more homogeneous in composition at the family (Myristicaceae, Arecaceae, Moraceae) and species levels (Condit 2002; Pitman *et al*. 2001), while species from eastern Amazonian have broadly different patterns of family-level composition being dominated by the Sapotaceae, Chrysobalanaceae, Fabaceae and Lecythidaceae (Chave *et al*. 2006). Wood density is driven by shifts in tree species composition (Terborgh & Andresen 1998) and tends to peak in the slow-growing forests on infertile soils in eastern Amazon and the Guyanas (Malhi *et al*. 2006). Soil physical properties in combination with limited nutrient supply in eastern Amazon favour slow-growing species and increases species that invest their resources in structures that can support taller and bigger trees with a long lifespan (Malhi *et al*. 2004; Quesada *et al*. 2009). Temperature and dry season precipitation effects on the structure and wood density are more ambiguous (Quesada *et al*. 2012), although species with higher wood density are better able to resist drought-induced embolism (Hacke *et al*. 2001) and therefore tolerate longer periods of high vapor water deficit and evaporative demand (McDowell *et al*. 2018). This myriad of environmental variables with confounding effects on species composition, as well as on their physiological and structural traits, play a crucial role in the tree lifespan and the size of trees (Muller-Landau 2004).

### Conditions supporting tall trees

In our study, the low wind speed was determined as the single most important predictor of the occurrence of the tallest trees in the Brazilian Amazon. The fact that trees adapt to their wind environment and are shorter in windy locations has been widely observed in temperate regions (Bonnesoeur *et al*. 2016; Telewski 2006). We can see a similar effect across the Amazon, with trees over 70 m tall having a 50-75% likelihood of surviving in the calmest areas but a sharply decreasing probability with stronger winds. This agrees with previous findings that disturbance rates are far higher in the Western Amazon (Espírito-Santo *et al*. 2014) and may demonstrate how significant the role of wind is in shaping the niche for extraordinarily tall trees. The importance of wind speed was also apparent in the Random Forest model which showed a 9 m reduction in the estimated tree height from the calmest to the windiest areas (Figure 2). The zonal velocity (i.e. the eastward component), which is the prevailing wind direction in the region, drives this pattern. Interestingly, our data showed that the lightning rate was only weakly related to maximum forest height patterns in both the Random Forests and MaxEnt models. Despite being relevant to the death of individual trees (Marra *et al*. 2014; Bonnesoeur *et al*. 2016; Niklas 1998) and being the key factor causing tree deaths in tropical forests of Panama (Yanoviak *et al*. 2019), lightning and storms do not seem to impact the potential dominant tree of a region, nor to limit the survival of the tallest trees, in light of our results.

A balance between tree structural strength and wind shearing forces contributes to set an upper limit to tree height development (Klein *et al*. 2015). The wind has a direct effect on tree height, since trees adapt their growth rates to their local wind environment, although the scale of this effect is unknown (Telewski 2006; Bonnesoeur *et al*. 2016). Extreme wind speeds, often associated with convective storms in the tropics, can also snap or uproot trees. Large-scale wind patterns in the Amazon are dominated by the easterly trade winds. Wind damage is most common from September to February (Negrón-Juárez *et al*. 2017) and taller trees have higher rates of mortality in wind storms (Rifai *et al*. 2016). Remote sensing analyses have shown that disturbance rates are much higher in the western Amazon compared to the east (Espírito-Santo *et al*. 2014).

A decrease in cloud-free days goes together with an increase in solar radiation (Barkhordarian *et al*. 2019), which, along with changes in the Vapor Pressure Deficit, or atmospheric dryness, drive changes in the physiological function of trees (Williams *et al*. 2012; Nunes *et al*. 2019). The increase in diffuse radiation led by cloudy conditions induces an increase in photosynthetic activity (Gu 2003). Tree responses to direct solar radiation are dependent on the species and developmental stage, with physiological and structural changes to maximize either growth or survival (Wright *et al*. 2004; Nunes *et al*. 2019; Poorter & Bongers 2006). As the traits of individual trees are at least conserved at the species level, additional variation is determined by the local environment (Fyllas *et al*. 2009). As trees grow taller, increasing leaf water stress due to gravity and path length resistance may limit leaf expansion and photosynthesis for further height growth (Koch *et al*. 2004). Tall trees have direct exposure to sunlight and high temperatures lead to higher stomatal control to avoid excessive water loss (Drake *et al*. 2018; Rowland *et al*. 2015).

Elevation was also a key predictor of tree height, with low-lying forests growing potentially less than trees in terrains over 40 m a.s.l. (Fig. 4). The topographic gradient is likely to be related to the likelihood of flooding in the low elevation transects on the lowlands. Rivers erode the *terra firme* terraces and create floodplains of variable sizes dating to the Miocene, with terrace–floodplain elevation differences decreasing eastwards from the Andes (Hamilton *et al*. 2007). Shifts in multiple canopy chemical traits between the terrace and floodplain forests in the Amazon are paralleled by species turnover, which reveals the micro-topography effects on the growth-defense trade-off in Amazonian forests, and its associated processes of nutrient mobilization and deposition (Asner *et al*. 2015). The species and trait shifts with topographical variation in the Amazon also confers an adaptive drought resistance, with species from the plateaus more susceptible to prolonged periods with lower soil water content, and, therefore, investing in higher hydraulic safety with higher wood density, lower mean vessel hydraulic diameter, lower mean vessel area and smaller stem cross-sectional sapwood area than species in valley forests (Cosme *et al*. 2017).

An increase in soil clay content also translated into an increase in maximum height. Clay content is usually highest on flat terrain (Laurance *et al*. 1999) decreasing from about 75% to 5% when moving from the plateau areas to the valleys (Ferraz *et al*. 1998; Toledo *et al*. 2016). Previous studies also indicated the presence of clayey soil in the plateau areas of the Amazon (Broedel *et al. 2017*; Cerri & Volkoff 1987; Marques *et al*. 2002; Marques *et al*. 2004; Marques *et al*. 2015). Our results suggest that 1) if the clayey soils of our study occur in the plateau areas with lower soil water content, a shift of species associated with the plateaus favoured species with higher hydraulic safety, otherwise 2) access to structured soils seems to be essential for trees to grow taller. A previous study showed an increase in wood density from stands on sandy soils in valleys to clayey soils on plateaus at a local scale in Central Amazon, and lower tree mortality rates in clayey soils (Toledo *et al*. 2016). These patterns were primarily driven by soil moisture - correlated to depth to water table - causing shifts in tree community composition (Schietti *et al*. 2013), and favouring higher hydraulic safety in the lower soil moisture areas of the plateaus (Toledo *et al*. 2016; Cosme *et al*. 2017). We suggest that the structured soils allow trees to obtain an additional volume of water during the dry season towards eastern Amazon, where soils tend to be richer in clay compared to central and western Amazon (Fisher *et al*. 2008; Hodnett *et al*. 1997). The dimorphic root systems associated with structured, clayey soils can redistribute water from deep layers to the soil surface during periods of drought (Broedel *et al*. 2017).

Chemical and physical properties of soils across the Amazon Basin tend to correlate with variations in and type of parent material, and exhibit an east-west soil age gradient (Quesada *et al*. 2011). This edaphic variation across geological formations has strong influences on the floristic, structural and demographic patterns in the Amazon (Quesada *et al*. 2012; ter Steege *et al*. 2006), with abrupt changes in species composition following changes in soil properties and topography (Phillips *et al*. 2003; Higgins *et al*. 2011). These patterns reflect more than a simple east-west gradient, due to a complex history of deposition and erosion dating to the Miocene (Higgins *et al*. 2011). Despite the clear heterogeneity caused by abrupt edaphic variation, two main gradients explain 24% of the total variation in tree community composition: one from the Guiana Shield to the southwestern Amazon, congruent with variation in soil fertility and its effects on tree wood density and seed mass, and another gradient from Colombia to the southeastern Amazon related to the length of the dry season (ter Steege *et al*. 2006). These gradients have distinctions in terms of their most abundant genera and occurrence of the Fabaceae family, which contains most of the large trees and grow successfully in low-dynamics environments such as the Guiana Shields. Higher occurrence of the Fabaceae in these low-fertility soils may occur due to the ability to fix nitrogen in the soil and ectomycorrhizal association (Webb & Sprent 2002; Sprent 2009).

Our results also demonstrate mean annual precipitation as a key factor for trees to grow taller. A tolerance curve associated the height of tall trees with precipitation, peaking at 2,300 mm yr^-1^ as optimal, but also showing that areas too dry or too wet may both inhibit the growth of tall trees. Thus, we observed 4 m decline in maximum tree height in regions with annual precipitations below 1,500 mm yr^-1^ or above 3,000 mm yr^-1^. The availability of soil water depends on both precipitation and evapotranspiration, and our results suggest that below 1,500 mm yr^-1^ evapotranspiration may exceed precipitation in the Amazon (Scheffer *et al*. 2018), and mortality by the hydraulic failure may occur for trees near their maximum height (McDowell *et al*. 2008). Mean annual precipitation above 2,300 mm year^-1^ may be related to exceeding water, and the combination of high precipitation and poorly drained soils may result in anaerobic conditions with negative effects on tree growth and survival (Quesada *et al*. 2009). Furthermore, higher precipitation tends to be related to the occurrence of storms and stronger winds with increases in tree mortality (Aleixo *et al*. 2019).

Temperature and precipitation are key variables modulating the composition of species in the north-western to southeastern seasonality gradient (ter Steege *et al*. 2006). The mean precipitation in the Brazilian Amazon varies from less than 2,000 mm year^-1^ (in the south, east, and extreme north) to more than 3,000 mm year^-1^ (in the northwest) (Liebmann & Marengo 2001). The annual convective movement of the inter-tropical convergence zone results in distinct wet and dry seasons (Marengo & Nobre 2001). However, the dry season in the Amazon basin varies from virtually nonexistent to periods reaching up to seven consecutive months with less than 100 mm month^-1^ of rain (Sombroek 2001). A global analysis provided evidence for the control of water availability over forest canopy height around the world, but the predictability between wet/dry indicates the involvement of additional limiting factors as temperature or radiation (Klein *et al*. 2015).

## Conclusion

Plant size distributions can be understood as the demographic consequence of size-dependent variation in growth and mortality in old-growth forests, and the mortality of large trees is independent of resource availability and competition (Coomes *et al*. 2003). Understanding the spatial distribution of maximum tree height in tropical forests and how it is associated with environmental conditions and tree functional traits is of fundamental importance. Emergent trees that reach their maximum height are responsible for a significant amount of the transpired water flux and the above-ground carbon storage. Trees which reach these extraordinary heights are rare and only a small proportion of species have the necessary adaptions to achieve this. However, these adaptations are not sufficient alone, and maximum tree height is strongly influenced by environmental conditions. We found that, across the Brazilian Amazon, the most important conditions were a lower number of clear sky days (reducing stress from direct sunlight), and soil clay content (improving water retention). Our second analysis emphasized the importance of disturbance, showing that the tallest trees are only found in places with low wind speed, allowing trees to grow for centuries without substantial damage.

Current climate models differ in their predictions of large-scale changes in wind patterns, although warmer temperatures will mean that the air can hold more moisture, which will likely make convective storms more intense. Whatever the change in environmental conditions, it is likely to occur faster than trees can adapt. Our results showed that precipitation and temperature have a lower importance than expected from previous studies. Nevertheless, changes in the precipitation and radiation regimes (strongly linked to the number of cloudy days) could reshape our forest biomes. Ultimately, the association between environmental conditions and mechanisms of natural selection, where some traits have some advantages in comparison to others influencing the survival of the most adaptable, are key to understanding the complexity of this process in a changing climate.

## Acknowledgements

Funding was provided by the Coordenação de Aperfeiçoamento de Pessoal de Nível Superior Brasil (CAPES; Finance Code 001); Conselho Nacional de Desenvolvimento Científico e Tecnológico (Processes 403297/2016-8 and 301661/2019-7); Amazon Fund (grant 14.2.0929.1); National Academy of Sciences and US Agency for International Development (grant AID-OAA-A-11-00012); Universidade Federal dos Vales do Jequitinhonha e Mucuri (UFVJM); Instituto Nacional de Pesquisas Espaciais (INPE);

D. Almeida was supported by the São Paulo Research Foundation (#2018/21338-3 and #2019/14697-0);

B. Gimenez, G. Spanner and N. Higuchi were supported by INCT-Madeiras da Amazônia and Next Generation Ecosystem Experiments-Tropics (NGEE-Tropics), as part of DOE’s Terrestrial Ecosystem Science Program – Contract No. DE-AC02-05CH11231;

T. Jackson and D. Coomes were supported by the UK Natural Environment Research Council grant NE/S010750/1;

M. Nunes was supported by the Academy of Finland (decision number 319905);

J. Rosette was supported by the Royal Society University Research Fellowship (URF\R\191014);

## Author contributions

EBG, TJ, DC, MK, NH, MHN, JO conceived of the idea. EBG, MA, GS, FSP, AZM developed the analysis and performed the computations. EBG, MHN, TJ, MK, DV, RV, NH, CRR, RC, DAA, JR, BG, JO verified the results, interpreted the results, and wrote the manuscript.

## References

Abatzoglou, J.T., Dobrowski, S.Z., Parks, S.A. & Hegewisch, K.C. (2018). TerraClimate a high-resolution global dataset of monthly climate and climatic water balance from 1958–2015. Scientific Data, 5.

Albergel, C., Dutra, E., Bonan, B., Zheng, Y., Munier, S., Balsamo, G., et al. (2019). Monitoring and Forecasting the Impact of the 2018 Summer Heatwave on Vegetation. Remote Sensing, 11, 520.

Albrecht, R.I., Goodman, S.J., Buechler, D.E., Blakeslee, R.J. & Christian, H.J. (2016). Where Are the Lightning Hotspots on Earth?. Bulletin of the American Meteorological Society, 97, 2051–2068.

Aleixo, I., Norris, D., Hemerik, L., Barbosa, A., Prata, E., Costa, F., et al. (2019). Amazonian rainforest tree mortality driven by climate and functional traits. Nature Climate Change, 9, 384–388.

Almeida, C.T., Galvão, L.S., Ometto, J.P.H.B., Jacon, A.D., Souza Pereira, F.R. de, Sato, L.Y., et al. (2019). Combining LiDAR and hyperspectral data for aboveground biomass modeling in the Brazilian Amazon using different regression algorithms. Remote Sensing of Environment, 232, 111323.

Almeida, D.R.A., Stark, S.C., Schietti, J., Camargo, J.L.C., Amazonas, N.T., Gorgens, E.B., et al. (2019). Persistent effects of fragmentation on tropical rainforest canopy structure after 20 yr of isolation. Ecological Applications, 29.

Anderegg, W.R.L., Klein, T., Bartlett, M., Sack, L., Pellegrini, A.F.A., Choat, B., et al. (2016). Meta-analysis reveals that hydraulic traits explain cross-species patterns of drought-induced tree mortality across the globe. Proceedings of the National Academy of Sciences, 113, 5024–5029.

Arsanjani, J.J., Vaz, E., Bakillah, M. & Mooney, P. (2014). Towards initiating OpenLandMap founded on citizens’ science: The current status of land use features of OpenStreetMap in Europe.

Asner, G.P. (2009). Tropical forest carbon assessment: integrating satellite and airborne mapping approaches. Environmental Research Letters, 4, 034009.

Asner, G.P., Anderson, C.B., Martin, R.E., Tupayachi, R., Knapp, D.E. & Sinca, F. (2015). Landscape biogeochemistry reflected in shifting distributions of chemical traits in the Amazon forest canopy. Nature Geoscience, 8, 567–573.

Asner, G.P., Powell, G.V.N., Mascaro, J., Knapp, D.E., Clark, J.K., Jacobson, J., et al. (2010). High-resolution forest carbon stocks and emissions in the Amazon. Proceedings of the National Academy of Sciences, 107, 16738–16742.

Bae, S., Levick, S.R., Heidrich, L., Magdon, P., Leutner, B.F., Wöllauer, S., et al. (2019). Radar vision in the mapping of forest biodiversity from space. Nature Communications, 10.

Baker, T.R., Phillips, O.L., Malhi, Y., Almeida, S., Arroyo, L., Fiore, A.D., et al. (2004). Variation in wood density determines spatial patterns inAmazonian forest biomass. Global Change Biology, 10, 545–562.

Barkhordarian, A., Saatchi, S.S., Behrangi, A., Loikith, P.C. & Mechoso, C.R. (2019). A Recent Systematic Increase in Vapor Pressure Deficit over Tropical South America. Scientific Reports, 9.

Bennett, A.C., McDowell, N.G., Allen, C.D. & Anderson-Teixeira, K.J. (2015). Larger trees suffer most during drought in forests worldwide. Nature Plants, 1.

Bisht, G. & Bras, R.L. (2010). Estimation of net radiation from the MODIS data under all sky conditions: Southern Great Plains case study. Remote Sensing of Environment, 114, 1522–1534.

Bonnesoeur, V., Constant, T., Moulia, B. & Fournier, M. (2016). Forest trees filter chronic wind-signals to acclimate to high winds. New Phytologist, 210, 850–860.

Broedel, E., Tomasella, J., Cândido, L.A. & Randow, C. von. (2017). Deep soil water dynamics in an undisturbed primary forest in central Amazonia: Differences between normal years and the 2005 drought. Hydrological Processes, 31, 1749–1759.

Cao, M. & Woodward, F.I.N. (1998). Net primary and ecosystem production and carbon stocks of terrestrial ecosystems and their responses to climate change. Global Change Biology, 4, 185–198.

Cerri, C.C. & Volkoff, B. (1987). Carbon content in a yellow latosol of central Amazon rain forest.. ACTA OECOL.(OECOL. GEN.)., 8, 29–42.

Chave, J., Muller-Landau, H.C., Baker, T.R., Easdale, T.A., Steege, H.ter & Webb, C.O. (2006). Regional and phylogenetic variation of wood density across 2456 neotropical tree species. Ecological applications, 16, 2356–2367.

Chave, J., Piponiot, C., Maréchaux, I., de, F.H., Larpin, D., Fischer, F.J., et al. (2020). Slow rate of secondary forest carbon accumulation in the Guianas compared with the rest of the Neotropics.. Ecol Appl, 30, e02004.

Claverie, M., Matthews, J., Vermote, E. & Justice, C. (2016). A 30+ Year AVHRR LAI and FAPAR Climate Data Record: Algorithm Description and Validation. Remote Sensing, 8, 263.

Condit, R. (2002). Beta-Diversity in Tropical Forest Trees. Science, 295, 666–669.

Coomes, D.A., Dalponte, M., Jucker, T., Asner, G.P., Banin, L.F., Burslem, D.F.R.P., et al. (2017). Area-based vs tree-centric approaches to mapping forest carbon in Southeast Asian forests from airborne laser scanning data. Remote Sensing of Environment, 194, 77–88.

Coomes, D.A., Duncan, R.P., Allen, R.B. & Truscott, J. (2003). Disturbances prevent stem size-density distributions in natural forests from following scaling relationships. Ecology Letters, 6, 980–989.

Coomes, D.A., Jenkins, K.L. & Cole, L.E.S. (2006). Scaling of tree vascular transport systems along gradients of nutrient supply and altitude. Biology Letters, 3, 87–90.

Cosme, L.H.M., Schietti, J., Costa, F.R.C. & Oliveira, R.S. (2017). The importance of hydraulic architecture to the distribution patterns of trees in a central Amazonian forest. New Phytologist, 215, 113–125.

Dalponte, M. & Coomes, D.A. (2016). Tree-centric mapping of forest carbon density from airborne laser scanning and hyperspectral data. Methods in Ecology and Evolution, 7, 1236–1245.

Drake, J.E., Tjoelker, M.G., Vårhammar, A., Medlyn, B.E., Reich, P.B., Leigh, A., et al. (2018). Trees tolerate an extreme heatwave via sustained transpirational cooling and increased leaf thermal tolerance. Global Change Biology, 24, 2390–2402.

Diaz, S., Kattge, J., Cornelissen, J.H.C., Wright, I.J., Lavorel, S., Dray, S., et al. (2016). The global spectrum of plant form and function. Nature, 529, 167–171.

Enquist, B.J., Abraham, A.J., Harfoot, M.B.J., Malhi, Y. & Doughty, C.E. (2020). The megabiota are disproportionately important for biosphere functioning. Nature Communications, 11.

Espírito-Santo, F.D.B., Gloor, M., Keller, M., Malhi, Y., Saatchi, S., Nelson, B., et al. (2014). Size and frequency of natural forest disturbances and the Amazon forest carbon balance. Nature communications, 5, 1–6.

Farr, T.G., Rosen, P.A., Caro, E., Crippen, R., Duren, R., Hensley, S., et al. (2007). The shuttle radar topography mission. Reviews of geophysics, 45.

Fauset, S., Johnson, M.O., Gloor, M., Baker, T.R., Monteagudo, M.A., Brienen, R.J., et al. (2015). Hyperdominance in Amazonian forest carbon cycling.. Nat Commun, 6, 6857.

Feldpausch, T.R., Lloyd, J., Lewis, S.L., Brienen, R.J.W., Gloor, M., Monteagudo Mendoza, A., et al. (2012). Tree height integrated into pantropical forest biomass estimates. Biogeosciences, 3381–3403.

Ferraz, J., Ohta, S.A.L.L.E.S. & Sales, P.C.de. (1998). Distribuição dos solos ao longo de dois transectos em floresta primária ao norte de Manaus (AM). Higuchi, N., Campos, MAA, Sampaio, PTB, and dos Santos, J., Espaço Comunicaçao Ltda., Manaus, Brazil, 264.

Fick, S.E. & Hijmans, R.J. (2017). WorldClim 2: new 1-km spatial resolution climate surfaces for global land areas. International Journal of Climatology, 37, 4302–4315.

Fisher, R.A., Williams, M., Lourdes Ruivo, M. de, Costa, A.L. de & Meir, P. (2008). Evaluating climatic and soil water controls on evapotranspiration at two Amazonian rainforest sites. Agricultural and Forest Meteorology, 148, 850–861.

Funk, C., Peterson, P., Landsfeld, M., Pedreros, D., Verdin, J., Shukla, S., et al. (2015). The climate hazards infrared precipitation with stations—a new environmental record for monitoring extremes. Scientific Data, 2.

Fyllas, N.M., Patino, S., Baker, T.R., Nardoto, G.B., Martinelli, L.A., Quesada, C.A., et al. (2009). Basin-wide variations in foliar properties of Amazonian forest: phylogeny soils and climate. Biogeosciences, 6, 2677–2708.

Fyllas, N.M., Quesada, C.A. & Lloyd, J. (2012). Deriving Plant Functional Types for Amazonian forests for use in vegetation dynamics models. Perspectives in Plant Ecology Evolution and Systematics, 14, 97–110.

Givnish, T.J., Wong, S.C., Stuart-Williams, H., Holloway-Phillips, M. & Farquhar, G.D. (2014). Determinants of maximum tree height inEucalyptusspecies along a rainfall gradient in Victoria Australia. Ecology, 95, 2991–3007.

Gorgens, E.B., Motta, A.Z., Assis, M., Nunes, M.H., Jackson, T., Coomes, D., et al. (2019). The giant trees of the Amazon basin. Frontiers in Ecology and the Environment, 17, 373–374.

Greenwood, S., Ruiz-Benito, P., Martínez-Vilalta, J., Lloret, F., Kitzberger, T., Allen, C.D., et al. (2017). Tree mortality across biomes is promoted by drought intensity lower wood density and higher specific leaf area. Ecology Letters, 20, 539–553.

Gu, L. (2003). Response of a Deciduous Forest to the Mount Pinatubo Eruption: Enhanced Photosynthesis. Science, 299, 2035–2038.

Gorgens, E.B., Soares, C.P.B., Nunes, M.H. & Rodriguez, L.C.E. (2016). Characterization of Brazilian forest types utilizing canopy height profiles derived from airborne laser scanning. Applied Vegetation Science, 19, 518–527.

Hacke, U.G., Sperry, J.S., Pockman, W.T., Davis, S.D. & McCulloh, K.A. (2001). Trends in wood density and structure are linked to prevention of xylem implosion by negative pressure. Oecologia, 126, 457–461.

Hamilton, S.K., Kellndorfer, J., Lehner, B. & Tobler, M. (2007). Remote sensing of floodplain geomorphology as a surrogate for biodiversity in a tropical river system (Madre de Dios Peru). Geomorphology, 89, 23–38.

Higgins, M.A., Ruokolainen, K., Tuomisto, H., Llerena, N., Cardenas, G., Phillips, O.L., et al. (2011). Geological control of floristic composition in Amazonian forests. Journal of Biogeography, 38,2136–2149.

Hijmans, R.J., Cameron, S.E., Parra, J.L., Jones, P.G. & Jarvis, A. (2005). Very high resolution interpolated climate surfaces for global land areas. International Journal of Climatology, 25, 1965–1978.

Hodnett, M.G., Vendrame, I., Marques Filho, A.D.O., Oyama, M.D. & Tomasella, J. (1997). Soil water storage and groundwater behaviour in a catenary sequence beneath forest in central Amazonia: I. Comparisons between plateau, slope and valley floor. Hydrology and Earth System Sciences Discussions, 1.

Huete, A.R., Didan, K., Shimabukuro, Y.E., Ratana, P., Saleska, S.R., Hutyra, L.R., et al. (2006). Amazon rainforests green-up with sunlight in dry season. Geophysical Research Letters, 33.

Jagels, R., Equiza, M.A., Maguire, D.A. & Cirelli, D. (2018). Do tall tree species have higher relative stiffness than shorter species?. American Journal of Botany, 105, 1617–1630.

Kang, S., Running, S.W., Zhao, M., Kimball, J.S. & Glassy, J. (2005). Improving continuity of MODIS terrestrial photosynthesis products using an interpolation scheme for cloudy pixels. International Journal of Remote Sensing, 26, 1659–1676.

Klein, T., Randin, C. & Körner, C. (2015). Water availability predicts forest canopy height at the global scale. Ecology Letters, 18, 1311–1320.

Koch, G.W., Sillett, S.C., Jennings, G.M. & Davis, S.D. (2004). The limits to tree height. Nature, 428, 851–854.

Larjavaara, M. (2013). The world’s tallest trees grow in thermally similar climates. New Phytolo-gist, 202, 344–349.

Laurance, W.F., Fearnside, P.M., Laurance, S.G., Delamonica, P., Lovejoy, T.E., Merona, J.M.R.-de, et al. (1999). Relationship between soils and Amazon forest biomass: a landscape-scale study. Forest Ecology and Management, 118, 127–138.

Liebmann, B. & Marengo, J.A. (2001). Interannual variability of the rainy season and rainfall in the Brazilian Amazon Basin. Journal of Climate, 14, 4308–4318.

Lindenmayer, D.B. & Laurance, W.F. (2016). The Unique Challenges of Conserving Large Old Trees. Trends in Ecology & Evolution, 31, 416–418.

Liu, H., Gleason, S.M., Hao, G., Hua, L., He, P., Goldstein, G., et al. (2019). Hydraulic traits are coordinated with maximum plant height at the global scale. Science Advances, 5, eaav1332.

Liu, J.-kuan, Liu, D. & Alsdorf, D. (2014). Extracting Ground-Level DEM From SRTM DEM in Forest Environments Based on Mathematical Morphology. IEEE Transactions on Geoscience and Remote Sensing, 52, 6333–6340.

Malhi, Y., Baker, T.R., Phillips, O.L., Almeida, S., Alvarez, E., Arroyo, L., et al. (2004). The above-ground coarse wood productivity of 104 Neotropical forest plots. Global Change Biology, 10, 563–591.

Malhi, Y., Wood, D., Baker, T.R., Wright, J., Phillips, O.L., Cochrane, T., et al. (2006). The regional variation of aboveground live biomass in old-growth Amazonian forests. Global Change Biology, 12, 1107–1138.

Marengo, J.A. & Nobre, C. (2001). General Characteristics and variability of Climate in the Amazon Basin and its Links to the Global Climate System. In: The hydroclimatological framework of Amazonia, Biogeochemistry of Amazonia. Cambridge University Press.

Marques, J.D.de O., Libardi, P.L., Teixeira, W.G. & Reis, A.M. (2004). Estudo de parâmetros físicos, químicos e hidricos de um Latossolo Amarelo, na regiâo Amazônica. Acta amazônica, 34, 145–154.

Marques, J.D.de O., Luizão, F.J., Teixeira, W.G., Sarrazin, M., Ferreira, S.J.F., Beldini, T.P., et al. (2015). Distribution of organic carbon in different soil fractions in ecosystems of central Amazonia. Revista Brasileira de Ciência do Solo, 39, 232–242.

Marques, J.J., Teixeira, W.G., Schulze, D.G. & Curi, N. (2002). Mineralogy of soils with unusually high exchangeable Al from the western Amazon Region. Clay Minerals, 37, 651–661.

Marra, D.M., Chambers, J.Q., Higuchi, N., Trumbore, S.E., Ribeiro, G.H.P.M., Santos, J. dos, et al. (2014). Large-Scale Wind Disturbances Promote Tree Diversity in a Central Amazon Forest. PLoS ONE, 9, e103711.

Marvin, D.C., Asner, G.P., Knapp, D.E., Anderson, C.B., Martin, R.E., Sinca, F., et al. (2014). Amazonian landscapes and the bias in field studies of forest structure and biomass. Proceedings of the National Academy of Sciences, 111, E5224–E5232.

Mason, P.J., Zillman, J.W., Simmons, A., Lindstrom, E.J., Harrison, D.E., Dolman, H., et al. (2010). Implementation plan for the global observing system for climate in support of the UN-FCCC (2010 Update).

McDowell, N., Allen, C.D., Anderson-Teixeira, K., Brando, P., Brienen, R., Chambers, J., et al. (2018). Drivers and mechanisms of tree mortality in moist tropical forests. New Phytologist, 219, 851–869.

McDowell, N., Pockman, W.T., Allen, C.D., Breshears, D.D., Cobb, N., Kolb, T., et al. (2008). Mechanisms of plant survival and mortality during drought: why do some plants survive while others succumb to drought?. New Phytologist, 178, 719–739.

McDowell, N.G. & Allen, C.D. (2015). Darcy’s law predicts widespread forest mortality under climate warming. Nature Climate Change, 5, 669–672.

Moles, A.T., Warton, D.I., Warman, L., Swenson, N.G., Laffan, S.W., Zanne, A.E., et al. (2009). Global patterns in plant height. Journal of Ecology, 97, 923–932.

Morrone, J.J. (2014). Biogeographical regionalisation of the Neotropical region. Zootaxa, 3782, 1.

Morton, D.C., Nagol, J., Carabajal, C.C., Rosette, J., Palace, M., Cook, B.D., et al. (2014). Amazon forests maintain consistent canopy structure and greenness during the dry season. Nature, 506, 221–224.

Muller-Landau, H.C. (2004). Interspecific and Inter-site Variation in Wood Specific Gravity of Tropical Trees. Biotropica, 36, 20–32.

Negron-Juarez, R.I., Jenkins, H.S., Raupp, C.F.M., Riley, W.J., Kueppers, L.M., Magnabosco Marra, D., et al. (2017). Windthrow Variability in Central Amazonia. Atmosphere, 8.

Niklas, K.J. (2007). Maximum plant height and the biophysical factors that limit it. Tree Physiology, 27, 433–440.

Niklas, K.J. (1998). The influence of gravity and wind on land plant evolution. Review of Palaeobotany and Palynology, 102, 1–14.

Nunes, M.H., Both, S., Bongalov, B., Brelsford, C., Khoury, S., Burslem, D.F.R.P., et al. (2019). Changes in leaf functional traits of rainforest canopy trees associated with an El Niño event in Borneo. Environmental Research Letters, 14, 085005.

Olauson, J. (2018). ERA5: The new champion of wind power modelling?. Renewable Energy, 126, 322–331.

Patino, S., Fyllas, N.M., Baker, T.R., Paiva, R., Quesada, C.A., Santos, A.J.B., et al. (2012). Coordination of physiological and structural traits in Amazon forest trees. Biogeosciences, 9, 775–801.

Pennisi, E. (2019). Forest giants are the trees most at risk. Science, 365, 962–963.

Pereira, I., Nascimento, H.M. do, Vicari, M.B., Disney, M., DeLucia, E., Domingues, T., et al. (2019). Performance of Laser-Based Electronic Devices for Structural Analysis of Amazonian Terra-Firme Forests. Remote Sensing, 11, 510.

Phillips, O.L., Baker, T.R., Arroyo, L., Higuchi, N., Killeen, T.J., Laurance, W.F., et al. (2004). Pattern and process in Amazon tree turnover, 1976-2001. Philosophical Transactions of the Royal Society of London. Series B: Biological Sciences, 359, 381–407.

Phillips, O.L., Vargas, P.N., Monteagudo, A.L., Cruz, A.P., Zans, M.-E.C., Sánchez, W.G., et al. (2003). Habitat association among Amazonian tree species: a landscape-scale approach. Journal of Ecology, 91, 757–775.

Phillips, S.J., Anderson, R.P. & Schapire, R.E. (2006). Maximum entropy modeling of species geographic distributions. Ecological Modelling, 190, 231–259.

Pitman, N.C.A., Terborgh, J.W., Silman, M.R., Núñez V, P., Neill, D.A., Cerón, C.E., et al. (2001). Dominance and distribution of tree species in upper Amazonian terra firme forests. Ecology, 82, 2101–2117.

Poorter, L. & Bongers, F. (2006). Leaf traits are good predictors of plant performance across 53 rain forest species. Ecology, 87, 1733–1743.

Powers, J.S., Vargas-G, G., Brodribb, T.J., Schwartz, N.B., Perez-Aviles, D., Smith-Martin, C.M., et al. (2020). A catastrophic tropical drought kills hydraulically vulnerable tree species. Global Change Biology.

Quesada, C.A., Lloyd, J., Anderson, L.O., Fyllas, N.M., Schwarz, M. & Czimczik, C.I. (2011). Soils of Amazonia with particular reference to the RAINFOR sites. Biogeosciences, 8,1415–1440.

Quesada, C.A., Lloyd, J., Schwarz, M., Baker, T.R., Phillips, O.L., Patiño, S., et al. (2009). Regional and large-scale patterns in Amazon forest structure and function are mediated by variations in soil physical and chemical properties. Biogeosciences Discussion, 6, 3993–4057.

Quesada, C.A., Phillips, O.L., Schwarz, M., Czimczik, C.I., Baker, T.R., Patiño, S., et al. (2012). Basin-wide variations in Amazon forest structure and function are mediated by both soils and climate. Biogeosciences, 9, 2203–2246.

Ramon, J., Lledó, L., Torralba, V., Soret, A. & Doblas-Reyes, F.J. (2019). What global reanalysis best represents near-surface winds?. Quarterly Journal of the Royal Meteorological Society, 145, 3236–3251.

Rifai, S.W., Urquiza Muñoz, J.D., Negrón-Juárez, R.I., Ramírez Arévalo, F.R., Tello-Espinoza, R., Vanderwel, M.C., et al. (2016). Landscape-scale consequences of differential tree mortality from catastrophic wind disturbance in the Amazon. Ecological Applications, 26, 2225–2237.

Rowland, L., Costa, A.C.L. da, Galbraith, D.R., Oliveira, R.S., Binks, O.J., Oliveira, A.A.R., et al. (2015). Death from drought in tropical forests is triggered by hydraulics not carbon starvation. Nature, 528, 119–122.

Rueda, M., Godoy, O. & Hawkins, B.A. (2016). Spatial and evolutionary parallelism between shade and drought tolerance explains the distributions of conifers in the conterminous United States. Global Ecology and Biogeography, 26, 31–42.

Ruhoff, A.L., Paz, A.R., Collischonn, W., Aragao, L.E.O.C., Rocha, H.R. & Malhi, Y.S. (2012). A MODIS-Based Energy Balance to Estimate Evapotranspiration for Clear-Sky Days in Brazilian Tropical Savannas. Remote Sensing, 4, 703–725.

Sakschewski, B., Bloh, W. von, Boit, A., Poorter, L., Peña-Claros, M., Heinke, J., et al. (2016). Resilience of Amazon forests emerges from plant trait diversity. Nature Climate Change, 6, 1032–1036.

Santos, E.G.D., Shimabukuro, Y.E., Moura, Y.M.D., Gonçalves, F.G., Jorge, A., Gasparini, K.A., et al. (2019). Multi-scale approach to estimating aboveground biomass in the Brazilian Amazon using Landsat and LiDAR data. International Journal of Remote Sensing, 40, 8635–8645.

Scheffer, M., Xu, C., Hantson, S., Holmgren, M., Los, S.O. & Nes, E.H. van. (2018). A global climate niche for giant trees. Global Change Biology, 24, 2875–2883.

Schietti, J., Emilio, T., Rennó, C.D., Drucker, D.P., Costa, F.R.C., Nogueira, A., et al. (2013). Vertical distance from drainage drives floristic composition changes in an Amazonian rainforest. Plant Ecology & Diversity, 7, 241–253.

Schimel, D., Pavlick, R., Fisher, J.B., Asner, G.P., Saatchi, S., Townsend, P., et al. (2015). Observing terrestrial ecosystems and the carbon cycle from space. Global Change Biology, 21, 1762–1776.

Simard, M., Fatoyinbo, L., Smetanka, C., Rivera-Monroy, V.H., Castañeda-Moya, E., Thomas, N., et al. (2018). Mangrove canopy height globally related to precipitation temperature and cyclone frequency. Nature Geoscience, 12, 40–45.

Sombroek, W. (2001). Spatial and Temporal Patterns of Amazon Rainfall. AMBIO: A Journal of the Human Environment, 30, 388–396.

Sprent, J.I. (2009). Legume Nodulation. Wiley-Blackwell.

Stovall, A.E.L., Shugart, H. & Yang, X. (2019). Tree height explains mortality risk during an intense drought. Nature Communications, 10.

Stropp, J., Umbelino, B., Correia, R.A., Campos-Silva, J.V., Ladle, R.J. & Malhado, A.C.M. (2020). The ghosts of forests past and future: deforestation and botanical sampling in the Brazilian Amazon. Ecography.

Tao, X., Liang, S., He, T. & Jin, H. (2016). Estimation of fraction of absorbed photosynthetically active radiation from multiple satellite data: Model development and validation. Remote Sensing of Environment, 184, 539–557.

Tejada, G., Görgens, E.B., Espírito-Santo, F.D.B., Cantinho, R.Z. & Ometto, J.P. (2019). Evaluating spatial coverage of data on the aboveground biomass in undisturbed forests in the Brazilian Amazon. Carbon Balance and Management, 14.

Telewski, F.W. (2006). A unified hypothesis of mechanoperception in plants. American Journal of Botany, 93, 1466–1476.

Terborgh, J. & Andresen, E. (1998). The composition of Amazonian forests: patterns at local and regional scales. Journal of Tropical Ecology, 14, 645–664.

Toledo, J.J., Castilho, C.V., Magnusson, W.E. & Nascimento, H.E.M. (2016). Soil controls biomass and dynamics of an Amazonian forest through the shifting of species and traits. Brazilian Journal of Botany, 40, 451–461.

Tuomisto, H., doninck, J.V., Ruokolainen, K., Moulatlet, G.M., Figueiredo, F.O.G., Sirén, A., et al. (2019). Discovering floristic and geoecological gradients across Amazonia. Journal of Biogeography, 46, 1734–1748.

Webb, J.R. & Sprent, J.I. (2002). Nodulation in Legumes. Kew Bulletin, 57, 634.

Williams, A.P., Allen, C.D., Macalady, A.K., Griffin, D., Woodhouse, C.A., Meko, D.M., et al. (2012). Temperature as a potent driver of regional forest drought stress and tree mortality. Nature Climate Change, 3, 292–297.

Wright, I.J., Reich, P.B., Westoby, M., Ackerly, D.D., Baruch, Z., Bongers, F., et al. (2004). The worldwide leaf economics spectrum.. Nature, 428, 821–7.

Yang, Y., Saatchi, S., Xu, L., Yu, Y., Lefsky, M., White, L., et al. (2016). Abiotic Controls on Macroscale Variations of Humid Tropical Forest Height. Remote Sensing, 8, 494.

Yanoviak, S.P., Gora, E.M., Bitzer, P.M., Burchfield, J.C., Muller-Landau, H.C., Detto, M., et al. (2019). Lightning is a major cause of large tree mortality in a lowland neotropical forest. New Phytologist, 225, 1936–1944.

Steege, H. ter, Pitman, N.C.A., Phillips, O.L., Chave, J., Sabatier, D., Duque, A., et al. (2006). Continental-scale patterns of canopy tree composition and function across Amazonia. Nature, 443, 444–447.

Gelder, H.A. van, Poorter, L. & Sterck, F.J. (2006). Wood mechanics allometry, and life-history variation in a tropical rain forest tree community. New Phytologist, 171, 367–378.

